# Partitioning the temporal changes in abundance-based beta diversity into loss and gain components

**DOI:** 10.1101/2021.05.28.446095

**Authors:** Shinichi Tatsumi, Ryosuke Iritani, Marc W. Cadotte

## Abstract

1. Ecologists have long recognized that the losses and gains in local species abundances can either decrease or increase spatial beta diversity, phenomena often referred to as biotic homogenization and differentiation, respectively. However, quantifying such dynamic impacts of species abundances on beta diversity has remained a methodological challenge.
2. Here, we develop a numerical method to additively partition the temporal changes in beta diversity into distinct components that reflect the losses and gains in local species abundances. Our method is based on Ružička and Bray–Curtis indices and the normalized abundance-based Whittaker’s beta diversity. The temporal changes in these measures are partitioned into components that represent biotic homogenization and differentiation driven by abundance losses and gains at both species and community levels.
3. Application of the method to a Swedish fish community dataset revealed decreases in beta diversity between 1990 and 2018. The homogenization of fish communities was explained by gains, but not losses, in species abundances across sites. Species-level partitioning further showed that the homogenization was largely caused by the increased population sizes of a particular species in sites where it was already present.
4. The results highlight that our partitioning method effectively identifies local population and community processes embedded in regional biodiversity patterns. We believe that explicit analyses of the losses and gains in species abundances should bring deeper insights into the dynamics of beta diversity.

## 1 INTRODUCTION

Beta diversity, the variation in the identities and abundances of species among sites, is a fundamental facet of biodiversity (Whittaker, 1960; Koleff et al., 2003; Anderson et al., 2011). Beta diversity can be quantified in two ways, namely by using incidence-based (i.e., presence–absence-based) and abundances-based approaches (Chao et al., 2005; Legendre & Legendre, 2012; Baselga, 2013). The two approaches weigh rare and dominant species differently, and thus offer complementary insights into community structure (Anderson et al., 2011; Legendre & Legendre, 2012; Li et al., 2016). Extensions of existing methods for analyzing incidence-based beta diversity to account for abundance can provide a more comprehensive understanding of biodiversity (Baselga, 2013; Chao et al., 2014).

While the replacement of endemic species by cosmopolitan nonnative species has been a global concern (McKinney & Lockwood, 1999), we still have mixed evidence for the consequent changes in beta diversity over time (Olden et al., 2018). The temporal decreases and increases in beta diversity, referred to as biotic homogenization and differentiation, respectively, subsume complex processes of local population dynamics (McKinney & Lockwood, 1999; Olden & Poff, 2003; Socolar et al., 2016; Rosenblad & Sax, 2017; Tatsumi et al., 2020). Empirical and simulation studies have shown that incidence- and abundance-based approaches can result in contrasting signs and magnitudes of temporal changes in beta diversity (Cassey et al., 2008; Li et al., 2016; Petersen et al., 2021). To build a more rigorous evidence base for biotic homogenization and differentiation, we need a tool to disentangle the processes underlying beta diversity changes.

Species extinctions and colonizations (i.e., changes in presence–absence status) can alter beta diversity in multiple ways (Olden & Poff, 2003; Rosenblad & Sax, 2017; Tatsumi et al., 2020, 2021). Specifically, extinctions lead to biotic homogenization when rare, infrequent species become regionally extinct, but otherwise result in differentiation (Socolar et al., 2016; Rosenblad & Sax, 2017; Tatsumi et al., 2020). Colonizations cause homogenization when species new to the region become widespread or existing species increase their regional dominance, but drive differentiation when new species colonize a small number of sites (Socolar et al., 2016; Rosenblad & Sax, 2017; Tatsumi et al., 2020). Extinctions and colonizations can also mask each other when one increases beta diversity and the other decreases it (Tatsumi et al., 2020, 2021). In our previous study (Tatsumi et al., 2021), we proposed a numerical method to additively partition such impacts of extinction and colonization on spatial beta diversity as quantified by incidence-based measures, namely Jaccard and Sørensen indices and Whittaker’s beta diversity.

Here, we develop a new method to additively partition the impacts of abundance losses and gains on spatial beta diversity by extending our previous incidence-based method. Similar to species extinctions and colonizations (i.e., binary changes between presence and absence), quantitative decreases and increases in local species abundances can drive either homogenization or differentiation (Socolar et al., 2016). The new method that we propose here allows one to partition such temporal changes in spatial variation (Δβ = β′ − β, where β and β′ are the values at *t* = 1 and 2, respectively) into distinct terms that reflect abundance losses and gains (Figure 1). Our method helps to resolve the local population dynamics and metacommunity processes embedded in regional biodiversity patterns using both incidence and abundance data.

**Figure 1.**
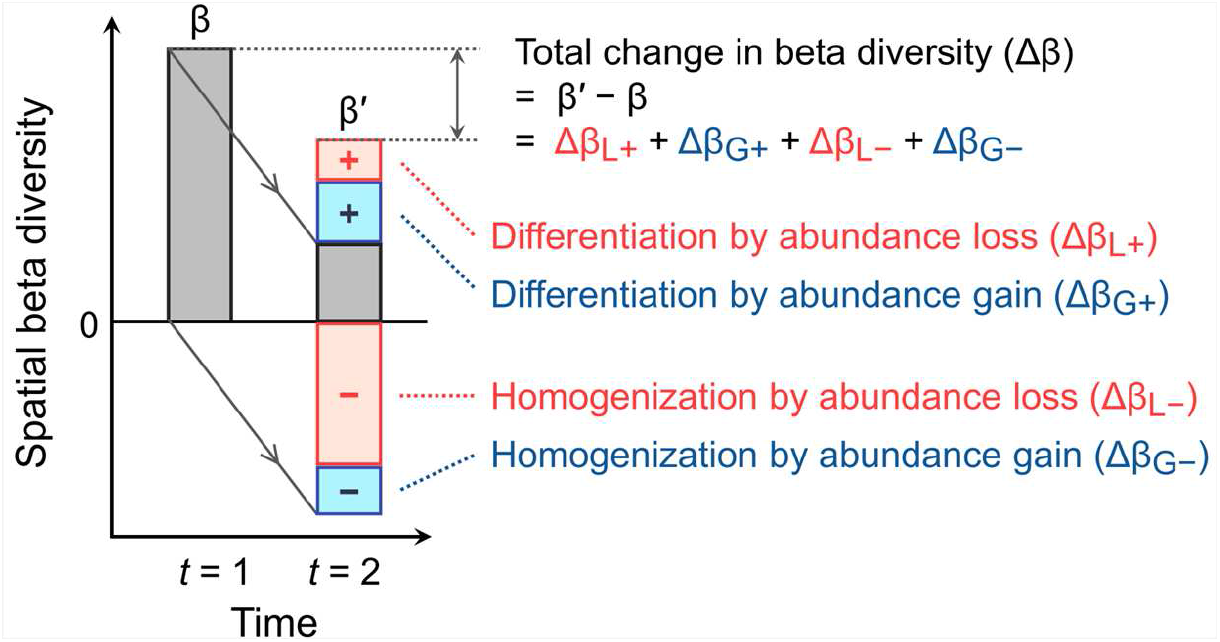
Temporal change in spatial beta diversity and its additive components. The components represent biotic homogenization or differentiation driven by losses or gains in species abundances.

## 2 METHODS

The partitioning method described below can be implemented using the ecopart R package available from GihHub: remotes::install_github(“communityecologist/ecopart”).

### 2.1 Partitioning equations

We describe the additive partitioning of temporal changes in pairwise dissimilarity measures, namely using Ružička (β_Ruž_) and Bray–Curtis indices (β_BC_) (Bray & Curtis, 1957; Ružička, 1958). These measures are defined as 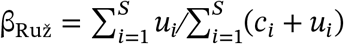 and 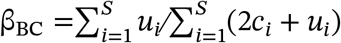, where *c*_*i*_ is the component of species abundance common to both sites, *u*_*i*_ is the component of species abundance unique to either site, *i* is species identity, and *S* is the number of species. Here, let 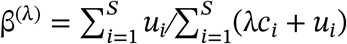, then β_Ruž_ = β^(1)^ and β_BC_ = β^(2)^. Ružička and Bray–Curtis indices are abundance-based extensions of Jaccard and Sørensen indices, respectively, for which the partitioning methods have already been described (Tatsumi et al., 2021). Although the Ružička and Bray–Curtis indices have different statistical properties from other common dissimilarity measures (e.g., Shannon entropy; Jost, 2007), we use the two indices here on account of their mathematical simplicity and wide uses in ecology.

Previous studies categorized species extinction and colonization into six types based on their impacts on spatial dissimilarity (Rosenblad & Sax, 2017; Tatsumi et al., 2020, 2021). We here extend these definitions to account for species abundances (Figure 2). The first type is the reduction in *u*_*i*_ (type 1 in Figure 2); that is, for the abundance of a given species *i* in site *k* (*a*_*ik*_), a component that was unique to either site (*u*_*i*_) at time *t* = 1 becomes lost at time *t* = 2. Type 2 is the case where a component of *a*_*ik*_ that was common to both sites (*c*_*i*_) becomes lost by an equal amount in both sites. Type 3 refers to the loss in *c*_*i*_ in the site where *a*_*ik*_ was smaller than, or equal to, that in the other site, turning *c*_*i*_ into *u*_*i*_. Type 4 refers to the gain in *u*_*i*_ in the site where *a*_*ik*_ was larger than, or equal to, that in the other site. Type 5 is the case where *c*_*i*_ increases by an equal amount in both sites. Type 6 refers to the gain in *c*_*i*_ in the site where *a*_*ik*_ was smaller than the other site, turning *u*_*i*_ into *c*_*i*_. Types 1, 5, and 6 decrease β^(λ)^, leading to homogenization, whereas types 2, 3, and 4 increase β^(λ)^, leading to differentiation. We write 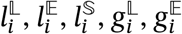, and 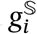 for the amount of changes in abundance that correspond to types 1, 2, 3, 4, 5, and 6, respectively (Appendix S1).

**Figure 2.**
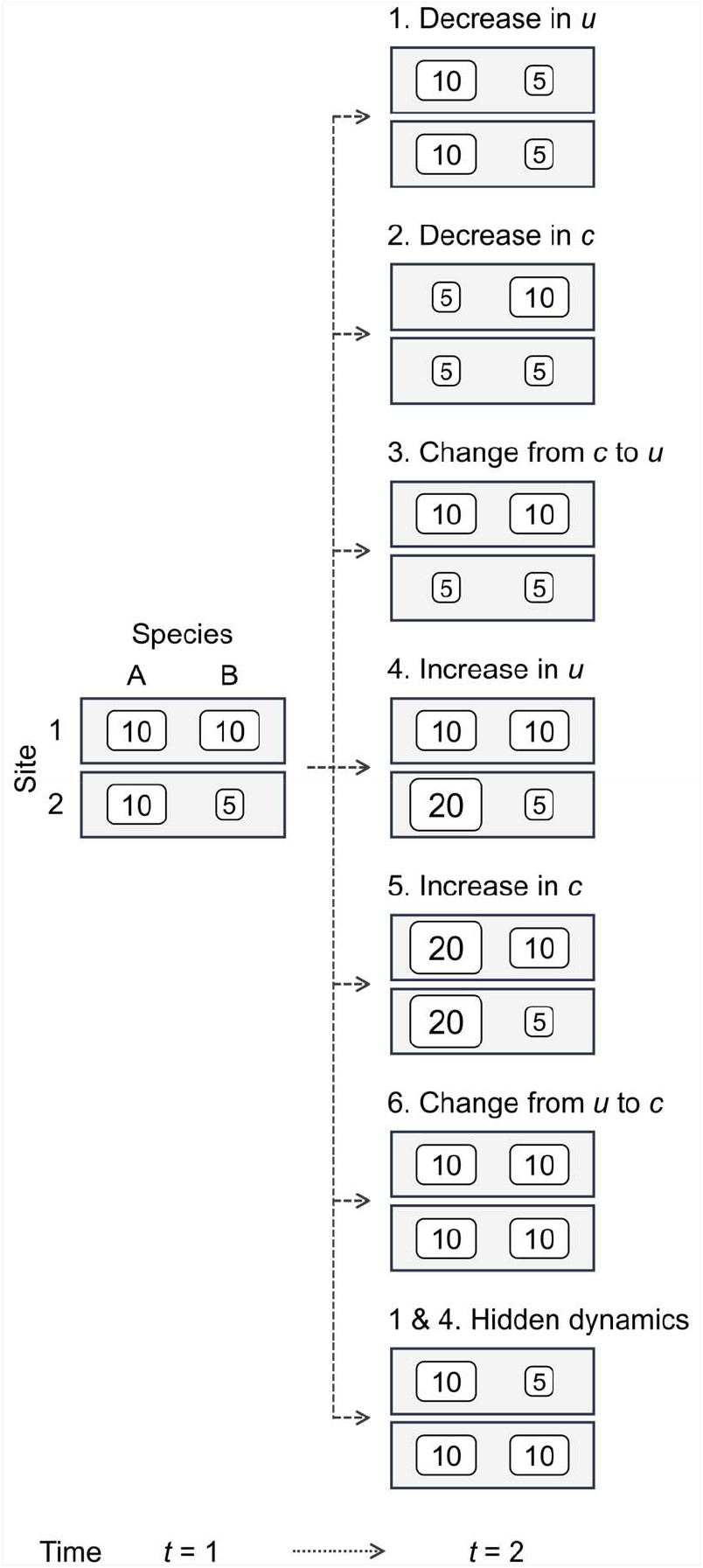
Schematic representation of six types of changes in beta diversity and hidden dynamics. The numbers in the boxes indicate species abundances. The variable *u* denotes the components of abundance unique to either site and *c* denotes the components of abundance common to both sites. For example, at *t* = 1, the *u* of species A is 0 and species B is 5. The *c* of species A is 10 and species B is 5. Thus, taking Ružička index as an example, β_Ruž_ = (0 + 5)/(10 + 5 + 0 + 5) = 1/4. In type 1 at *t* = 2, the *u* of species B becomes 0 as a result of an abundance loss in one of the two sites. Consequently, β_Ruž_ decreases to 0. In type 2, the *c* of species A is reduced to 5 and thus β_Ruž_ = (0 + 5)/(5 + 5 + 0 + 5) = 1/3. We can see from these examples that, while types 1 and 2 are both associated with abundance losses, the changes in β_Ruž_ can take either negative or positive values. It is such distinct types of changes in beta diversity our method allows one to partition.

It is possible for the abundance of a given species *i* to change differently in the two sites within the same time interval. Namely, if the abundance *a*_*ik*_ decreases in one site (e.g., *k* = 1) where it was larger and increases in the other site (e.g., *k* = 2) where it was smaller, then beta diversity can potentially show no net change (see the bottom case in Figure 2). We refer to such offsetting replacements in species abundances as hidden dynamics (Tatsumi et al. 2021). We write *d*_*i*_ for the changes in abundance that fall under this definition. In our partitioning method, we explicitly describe *d*_*i*_ as a distinct form of abundance losses and gains. In total, there are 32 possible ways *a*_*ik*_ can decrease and/or increase, including the hidden dynamics (Appendix S1). Further, beta diversity can be much more dynamic than any index portrays since abundances can change multiple times and in multiple ways between sampling intervals, and so here *c*_*i*_ and *u*_*i*_ refer to the net change between *t* = 1 and 2.

For brevity, we write the sum of a given variable across all species (1, 2, …, *S*) by using the uppercase letters (e.g., 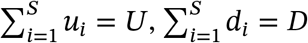, and 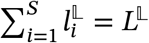). The temporal changes in *C* and *U* can then be written as Δ*C* = *C*′ − *C* = −*L*^𝔼^ − *L*^𝕊^ + *G*^𝔼^ + *G*^𝕊^ and Δ*U* = *U*′ − *U* = −*L*^𝕃^ + *L*^𝕊^ + *G*^𝕃^ − *G*^𝕊^. We can additively partition the temporal changes in pairwise dissimilarity (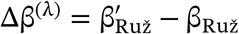 when λ = 1 and 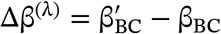 when λ = 2) into six terms that correspond to the six types of abundance changes:

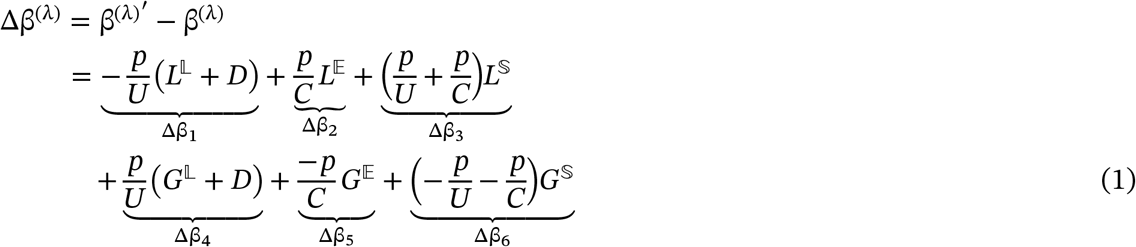

Where 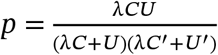 and *C* and *U* are both non-zero. See Appendix S1 for derivation of the equation.

The variable *D* denotes the hidden dynamics. The quantities 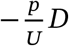 in Δβ_1_ and 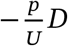 in Δβ_4_ cancel each other out by summing up to zero. These two quantities thus allow us, without causing any effect on Δβ, to explicitly account for *D* in a manner comparable to 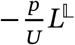 and 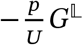.

The first three terms are associated with gains, while the last three are associated with losses. The six terms respectively correspond to the six types of abundance changes described in Figure 2. The first term, which is always negative, represents homogenization by abundance losses (Δβ_L−_). The second and third terms, which are always positive, indicate differentiation by abundance losses (Δβ_L+_). Similarly, the fourth term represents differentiation by abundance gains (Δβ_G+_) and the fifth and sixth terms indicate homogenization by abundance gains (Δβ_G−_). Depending on the ecological question at hand, one can sum the terms as Δβ_L_ = Δβ_L−_ + Δβ_L+_ and Δβ_G_ = Δβ_G−_ + Δβ_G+_ to assess the total impact of abundance losses and gains on Δβ, respectively (Figure 1).

See Appendix S2 for the responses of Δβ components (Δβ_L−_, Δβ_L+_, Δβ_G−_, and Δβ_G+_) to the abundance losses and gains (*L*^𝕃^, *L*^𝔼^, *L*^𝕊^, *G*^𝕃^, *G*^𝔼^, *G*^𝕊^, and *D*).

### 2.2 Multisite variation

Our partitioning method is applicable to multisite measures of beta diversity. Multisite measures are used to quantify variation among more than two sites (Baselga, 2010). Averaging pairwise dissimilarities (such as β_Ruž_ or β_BC_) across pairs of sites is a suboptimal approach due to their lack of statistical independence (Baselga, 2010, 2017).

In Appendix S3, we demonstrate the partitioning of multisite beta diversity by taking, as an example, the normalized abundance-based Whittaker’s beta diversity (β_W_) (cf. Baselga, 2017). We chose β_W_ here on account of its simple mathematical structure. Note, however, that future works are needed for partitioning other multisite indices, typically beta diversity based on Hill numbers (see *Discussion* for detail).

### 2.3 Species-level impacts on beta diversity

We can further use the partitioning equations to quantify the response of beta diversity to the losses and gains in the abundance of each species independently. For example, consider a case where the abundance of a species that had existed in one site was completely lost (i.e., the species went locally extinct). This loss will add 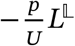 to Δβ in Eqn. 1. Thus, the added value can be interpreted as the consequence of the focal species’ abundance loss on beta diversity. In this way, Δβ can be additively partitioned into components that reflect the decreases and increases in the population size of individual species. Note that it is possible for the population size of a given species to decrease in some sites while increase in other sites within the same time interval, generating both the loss and gain components (i.e., Δβ_L−_, Δβ_L+_, Δβ_G−_, and Δβ_G+_).

## 3 APPLICATIONS

We applied the partitioning method to riverine fish community data retrieved from the Swedish Electrofishing Register database (Sers, 2013) via RivFishTIME (Comte et al., 2021). We used data collected in 65 waterbodies consisting of a total of 181 permanent sampling sites (2–10 sites per waterbody) across Sweden in 1990 and 2018 (see Appendix S4 for site IDs and selection criteria). The abundance of each fish species in each site was recorded as the number of individuals per 100 m^2^. We quantified the compositional variation among sites within each waterbody based on the normalized abundance-based Whittaker’s beta diversity using either incidence (presence–absence) or abundance data. The incidence data was obtained by transforming the abundance values larger than zero to one. We calculated the temporal changes in beta diversity (Δβ) and their additive components between 1990 and 2018. The components representing species extinctions and colonizations (i.e., changes from presence to absence and vice versa) are denoted as Δβ_E_ and Δβ_C_. Those that represent abundance losses and gains are denoted as Δβ_L_ and Δβ_G_.

The incidence- and abundance-based approaches provided complementary insights into the changes in beta diversity (Figure 3). While the incidence-based beta diversity showed no temporal trend (Figure 3a), the abundance-based beta diversity significantly decreased from 1990 and 2018, as indicated by Δβ less than zero (Figure 3b). The loss and gain components (Δβ_L_ and Δβ_G_) revealed that this homogenization of fish communities was explained by gains, but not losses, in species abundances (Figure 3b). Partitioning Δβ into species-level components further showed that the homogenization was largely caused by brown trout (*Salmo trutta*) (Figure 3d; see Appendix S4 for the results of all species). The fact that the colonization component Δβ_C_ of brown trout was not significant (Figure 3c) indicates that the homogenization did not result from colonizations of brown trout to new sites. Rather, it was caused by the increased sizes of brown trout populations (potentially associated with fishing restrictions and stocking; Almesjö & Limén, 2009) in sites where they were already present but in low abundances, leading to a more spatially-uniform abundance distribution.

**Figure 3.**
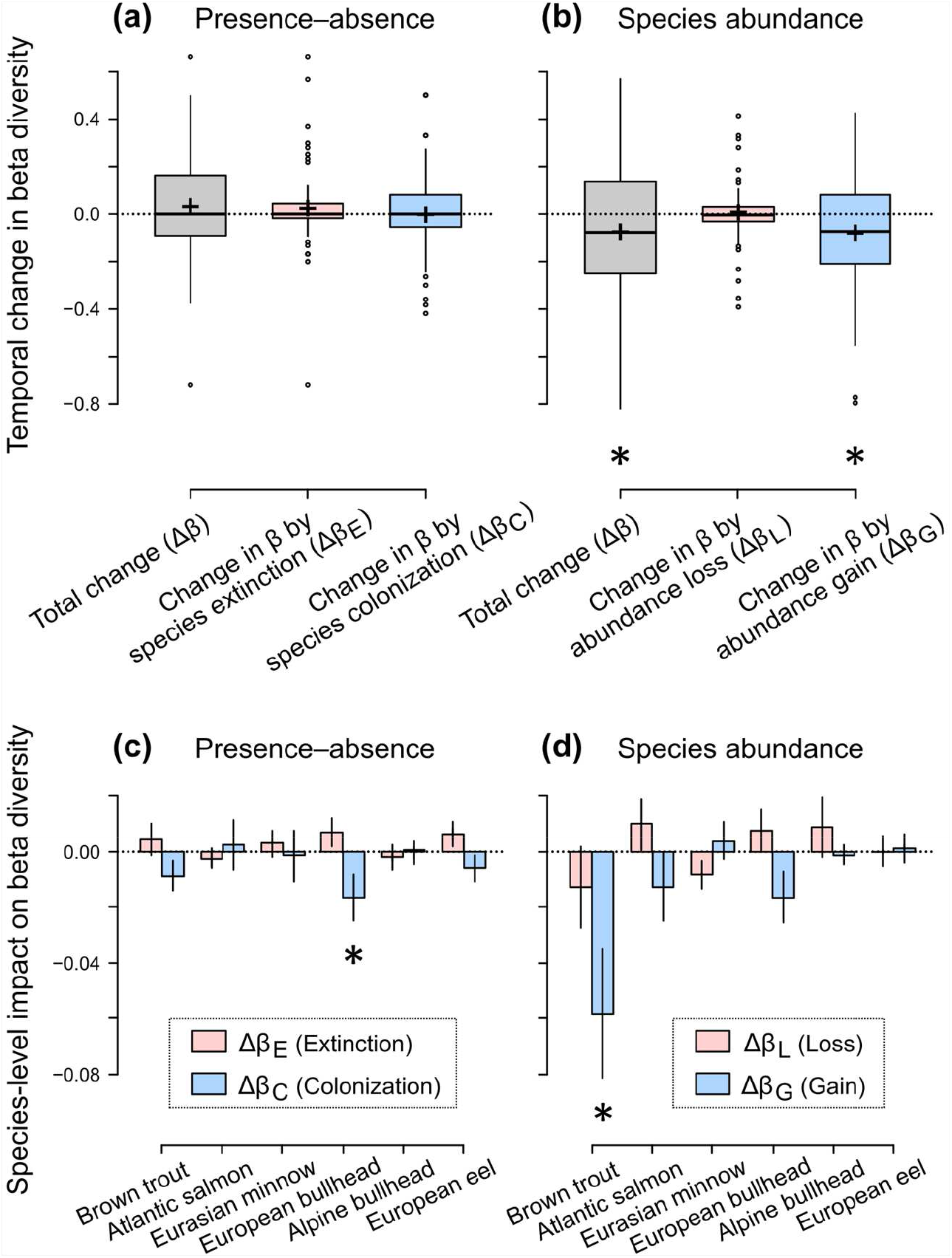
Temporal change in beta diversity and its components of riverine fish communities in 65 waterbodies across Sweden between 1990 and 2018. Beta diversity was defined as the compositional variation among multiple sampling sites within each waterbody. (a) Temporal changes in beta diversity based on species presence–absence and its extinction and colonization components. (b) Temporal changes in beta diversity based on species abundance and its loss and gain components. (c) Impacts of local extinctions and colonizations (i.e., changes from presence to absence and vice versa) of the six most abundant species (arranged in descending order) on beta diversity. (d) Impacts of abundance losses and gains of the six species on beta diversity. In the upper panels, the plus signs, horizontal lines, boxes, and circles indicate the means, medians, interquartile ranges (IQR), and outliers (values outside 1.5×IQR), respectively. In the bottom panels, bars and lines show the means ± standard errors. Asterisks indicate that the mean value was significantly different from zero (one-sample *t*-test; *P* < 0.05).

## 4 Discussion

We developed a new method to partition the temporal changes in beta diversity into distinct components that reflect the losses and gains in species abundances (Figure 1). The method provides a unified approach to analyze biotic homogenization and differentiation using both incidence and abundance data. Application of our method to an empirical dataset revealed different trends in incidence- and abundance-based beta diversity (Δβ_E_, Δβ_C_, Δβ_L_, and Δβ_G_) (Figure 3). The two approaches collectively showed that gains in abundances, but not colonization, of particular species made communities more similar over time (Figure 3). The results highlight that our partitioning method effectively identifies local population and community processes embedded in regional biodiversity patterns.

Moving forward, further generalizations of our temporal partitioning method are needed. In this study, we partitioned Ružička and Bray–Curtis indices and the normalized abundance-based Whittaker’s beta diversity. A promising next step will be to extend the method to beta diversity based on Hill numbers (Hill, 1973; Jost, 2007; Chao & Chiu, 2016). Hill numbers link different lines of beta diversity research together and unify multiple dissimilarity measures into a common expression (Jost, 2007; Chao & Chiu, 2016; Chao et al., 2019). Temporal partitioning of Hill-number-based beta diversity could thus give us a synthetic understanding of community dynamics. Exploring the potential connections between our method and other partitioning methods that are based on Hill numbers (Godsoe et al., 2021, 2022) would also be an important step forward.

We expect our partitioning method to serve as a useful tool in both basic and applied ecology. Specifically, the capability of our method to quantify species-level processes could help conservation practitioners to assess the impacts of particular species on regional biodiversity (e.g., increased abundance of an invasive nonnative species and consequent decreases in endangered species). Empirical ecologists could use the homogenization and differentiation components of beta diversity to infer metacommunity processes and regional coexistence mechanisms. We believe that explicit analyses of the losses and gains in species abundances bring deeper insights into ecological community structure across space and time.

## AUTHORS’ CONTRIBUTIONS

ST conceived the study, derived the partitioning equations, and analyzed the data. ST wrote the manuscript with inputs from RI and MWC. All authors contributed to manuscript revision.

## DATA ACCESSIBILITY

The R package ecopart (‘Ecological COmmunity PARTitioning’ or ‘Extinction and COlonization PARTitioning’) and an R script for extracting the Swedish fish community data from RivFishTIME (Comte et al., 2021) are available at GitHub (https://github.com/communityecologist). The ecopart package can be installed using the remotes package (Csardi et al., 2021): remotes::install_github(“communityecologist/ecopart”).

## ACKNOWLEDGEMENTS

We thank William Godsoe and an anonymous reviewer for constructive comments and people who contributed to the Swedish Electrofishing Register database (Sers, 2013). ST was supported by a JSPS Overseas Research Fellowship (No. 201860500) and a JSPS grant (No. 21K14880) from the Japan Society for the Promotion of Science. RI was supported by JSPS grants (No. 19K22457, 19K23768, and 20K15882).

## Notes

### Competing Interest Statement

The authors have declared no competing interest.

## REFERENCES

Almesjö, L., & Limén, H. (2009). Fish populations in Swedish waters: How are they influenced by fishing, eutrophication and contaminants? The Riksdag Printing Office.

Anderson, M. J., Crist, T. O., Chase, J. M., Vellend, M., Inouye, B. D., Freestone, A. L., … Swenson, N. G. (2011). Navigating the multiple meanings of beta diversity: A roadmap for the practicing ecologist. Ecology Letters, 14, 19–28.

Baselga, A. (2010). Partitioning the turnover and nestedness components of beta diversity. Global Ecology and Biogeography, 19, 134–143.

Baselga, A. (2013). Separating the two components of abundance-based dissimilarity: Balanced changes in abundance vs. abundance gradients. Methods in Ecology and Evolution, 4, 552–557.

Baselga, A. (2017). Partitioning abundance-based multiple-site dissimilarity into components: Balanced variation in abundance and abundance gradients. Methods in Ecology and Evolution, 8, 799–808.

Bray, J. R., & Curtis, J. T. (1957). An ordination of the upland forest communities of southern Wisconsin. Ecological Monographs, 27, 325–349.

Cassey, P., Lockwood, J. L., Olden, J. D., & Blackburn, T. M. (2008). The varying role of population abundance in structuring indices of biotic homogenization. Journal of Biogeography, 35, 884–892.

Chao, A., Chazdon, R. L., & Shen, T. J. (2005). A new statistical approach for assessing similarity of species composition with incidence and abundance data. Ecology Letters, 8, 148–159.

Chao, A., & Chiu, C. H. (2016). Bridging the variance and diversity decomposition approaches to beta diversity via similarity and differentiation measures. Methods in Ecology and Evolution, 7, 919–928.

Chao, A., Chiu, C., Wu, S., Huang, C., & Lin, Y. (2019). Comparing two classes of alpha diversities and their corresponding beta and (dis)similarity measures, with an application to the Formosan sika deer Cervus nippon taiouanus reintroduction programme. Methods in Ecology and Evolution, 10, 1286–1297.

Chao, A., Gotelli, N. J., Hsieh, T. C., Sander, E. L., Ma, K. H., Colwell, R. K., & Ellison, A. M. (2014). Rarefaction and extrapolation with Hill numbers: A framework for sampling and estimation in species diversity studies. Ecological Monographs, 84, 45–67.

Comte, L., Carvajal-Quintero, J., Tedesco, P. A., Giam, X., Brose, U., Erős, T., … Olden, J. D. (2021). RivFishTIME: A global database of fish time-series to study global change ecology in riverine systems. Global Ecology and Biogeography, 30, 38–50.

Csardi, G., Hester, J., Wickham, H., Chang, W., Morgan, M., & Tenenbaum, D. (2021). remotes: R package installation from remote repositories, including ‘GitHub’. R package version 2.4.2.

Godsoe, W., Bellingham, P. J., & Moltchanova, E. (2022). Disentangling niche theory and beta diversity change. The American Naturalist, 199, 510–522.

Godsoe, W., Eisen, K. E., Stanton, D., & Sirianni, K. M. (2021). Selection and biodiversity change. Theoretical Ecology, 14, 367–379.

Hill, M. O. (1973). Diversity and evenness: A unifying notation and its consequences. Ecology, 54, 427–432.

Jost, L. (2007). Partitioning diversity into independent alpha and beta components. Ecology, 88, 2427–2439.

Koleff, P., Gaston, K. J., & Lennon, J. J. (2003). Measuring beta diversity for presence-absence data. Journal of Animal Ecology, 72, 367–382.

Legendre, P., & Legendre, L. (2012). Numerical Ecology (3rd ed.). Elsevier.

Li, S., Cadotte, M. W., Meiners, S. J., Pu, Z., Fukami, T., & Jiang, L. (2016). Convergence and divergence in a long-term old-field succession: The importance of spatial scale and species abundance. Ecology Letters, 19, 1101–1109.

McKinney, M. L., & Lockwood, J. L. (1999). Biotic homogenization: A few winners replacing many losers in the next mass extinction. Trends in Ecology & Evolution, 14, 450–453.

Olden, J. D., Comte, L., & Giam, X. (2018). The Homogocene: A research prospectus for the study of biotic homogenisation. NeoBiota, 37, 23–36.

Olden, J. D., & Poff, N. L. (2003). Toward a mechanistic understanding and prediction of biotic homogenization. The American Naturalist, 162, 442–460.

Petersen, K. N., Freeman, M. C., Kirsch, J. E., McLarney, W. O., Scott, M. C., & Wenger, S. J. (2021). Mixed evidence for biotic homogenization of Southern Appalachian fish communities. Canadian Journal of Fisheries and Aquatic Sciences, 78, 1397–1406.

Rosenblad, K. C., & Sax, D. F. (2017). A new framework for investigating biotic homogenization and exploring future trajectories: Oceanic island plant and bird assemblages as a case study. Ecography, 40, 1040–1049.

Ružička, M. (1958). Anwendung mathematisch-statisticher Methoden in der Geobotanik (synthetische Bearbeitung von Aufnahmen). Biológia, Bratislava, 13, 647–661.

Sers, B. (2013). Swedish Electrofishing RegiSter – SERS. Swedish University of Agricultural Sciences (SLU), Department of Aquatic Resources. Retrieved from http://www.slu.se/electrofishingdatabase (December 21, 2021).

Socolar, J. B., Gilroy, J. J., Kunin, W. E., & Edwards, D. P. (2016). How should beta-diversity inform biodiversity conservation? Trends in Ecology & Evolution, 31, 67–80.

Tatsumi, S., Iritani, R., & Cadotte, M. W. (2021). Temporal changes in spatial variation: partitioning the extinction and colonisation components of beta diversity. Ecology Letters, 24, 1063–1072.

Tatsumi, S., Strengbom, J., Čugunovs, M., & Kouki, J. (2020). Partitioning the colonization and extinction components of beta diversity across disturbance gradients. Ecology, 101, e03183.

Whittaker, R. H. (1960). Vegetation of the Siskiyou Mountains, Oregon and California. Ecological Monographs, 30, 279–338.

