## Supporting Information for "Partitioning the temporal changes in abundance-based beta diversity into loss and gain components"

---

### Appendix S1: Derivation of partitioning equations: Pairwise dissimilarity

We first derive the equations to additively partition the temporal changes in Ružička and Bray–Curtis indices which are abundance-based, pairwise dissimilarity measures. We define the variables as follows:

| | |
|---|---|
| $\beta_{\text{Ruž}} = \frac{\sum_{i=1}^{S} u_i}{\sum_{i=1}^{S}(c_i+u_i)}$ | Ružička index. |
| $\beta_{\text{BC}} = \frac{\sum_{i=1}^{S} u_i}{\sum_{i=1}^{S}(2c_i+u_i)}$ | Bray–Curtis index. |
| $i$ | Species identity. |
| $S$ | Number of species. |
| $c_i$ | Component of species abundance common to both sites. |
| $u_i$ | Component of species abundance unique to either site. |
| $a_{ik}$ | Abundance of species $i$ in site $k$. |
| $l_i^{\mathbb{L}}$ | Type 1: Amount of loss in $u_i$ in the site where $a_{ik}$ was larger than that in the other site. |
| $l_i^{\mathbb{E}}$ | Type 2: An equal amount of loss in $c_i$ in both sites. |
| $l_i^{\mathbb{S}}$ | Type 3: Amount of loss in $c_i$ in the site where $a_{ik}$ was smaller than, or equal to, that in the other site. |
| $g_i^{\mathbb{L}}$ | Type 4: Amount of gain in $u_i$ in the site where $a_{ik}$ was larger than, or equal to, that in the other site. |
| $g_i^{\mathbb{E}}$ | Type 5: An equal amount of gain in $c_i$ in both sites. |
| $g_i^{\mathbb{S}}$ | Type 6: Amount of gain in $c_i$ in the site where $a_{ik}$ was smaller than that in the other site. |
| $d_i$ | Hidden dynamics: Amount of loss in $a_{ik}$ in the site where it was larger and gain in $a_{ik}$ in the other site where it was smaller. |

See also Figures S1 and S2 for visual descriptions of the variables.

For brevity, we write the sum of a given variable across all species (1, 2, ..., $S$) using the uppercase letters (e.g., $\sum_{i=1}^{S} c_i = C$ and $\sum_{i=1}^{S} l_i^{\mathbb{L}} = L^{\mathbb{L}}$). To distinguish the variables at $t = 1$ and $t = 2$, we add prime symbols to the latter cases (e.g. $U'$). From the definitions above, we can write

$$\Delta C = C' - C = -L^{\mathbb{E}} - L^{\mathbb{S}} + G^{\mathbb{E}} + G^{\mathbb{S}}$$
$$\Delta U = U' - U = -L^{\mathbb{L}} + L^{\mathbb{S}} + G^{\mathbb{L}} - G^{\mathbb{S}}.$$

The temporal changes in pairwise dissimilarity ($\Delta\beta^{(\lambda)} = \beta'_{\text{Ruž}} - \beta_{\text{Ruž}}$ when $\lambda = 1$, and $\beta^{(\lambda)} = \beta'_{\text{BC}} - \beta_{\text{BC}}$ when $\lambda = 2$) can then be written as

$$\begin{aligned}
\Delta\beta^{(\lambda)} &= \Delta\beta^{(\lambda)'} - \Delta\beta^{(\lambda)} \\
&= \frac{U'}{\lambda C' + U'} - \frac{U}{\lambda C + U} \\
&= \frac{\lambda(CU' - C'U)}{(\lambda C + U)(\lambda C' + U')} \\
&= \frac{\lambda CU\left(\frac{\Delta U}{U} - \frac{\Delta C}{C}\right)}{(\lambda C + U)(\lambda C' + U')} \qquad \because\; U' = U + \Delta U, C' = C + \Delta C \\
&= \underbrace{\frac{\lambda CU}{(\lambda C + U)(\lambda C' + U')}}_{p} \times \left(\frac{-L^{\mathbb{L}} + L^{\mathbb{S}} + G^{\mathbb{L}} - G^{\mathbb{S}}}{U} + \frac{L^{\mathbb{E}} + L^{\mathbb{S}} - G^{\mathbb{E}} - G^{\mathbb{S}}}{C}\right),
\end{aligned}$$

where $C$ and $U$ are both non-zero. Accordingly, the partitioning equation, without the hidden dynamics, reads

$$\begin{aligned}
\Delta\beta^{(\lambda)} &= \Delta\beta^{(\lambda)'} - \Delta\beta^{(\lambda)} \\
&= \underbrace{-\frac{p}{U}L^{\mathbb{L}}}_{\Delta\beta_1} + \underbrace{\frac{p}{C}L^{\mathbb{E}}}_{\Delta\beta_2} + \underbrace{\left(\frac{p}{U} + \frac{p}{C}\right)L^{\mathbb{S}}}_{\Delta\beta_3} + \underbrace{\frac{p}{U}G^{\mathbb{L}}}_{\Delta\beta_4} + \underbrace{\frac{-p}{C}G^{\mathbb{E}}}_{\Delta\beta_5} + \underbrace{\left(-\frac{p}{U} - \frac{p}{C}\right)G^{\mathbb{S}}}_{\Delta\beta_6}.
\end{aligned}$$

To account for hidden dynamics, we add $D$ to the multipliers of the first and fourth terms in the above equation. Hidden dynamics refer to decreases in species abundance in the site where it was larger and offsetting increases in species abundance in the other site where it was smaller (Figure S2). By adding $D$ to the equation, we get Equation 1 in the main text.

**(a)** Pairwise dissimilarity measures

Ružička

$$\beta_{\text{Ruž}} = \frac{\sum_{i=1}^{S} u_i}{\sum_{i=1}^{S} (c_i + u_i)}$$

Bray–Curtis

$$\beta_{\text{BC}} = \frac{\sum_{i=1}^{S} u_i}{\sum_{i=1}^{S} (2c_i + u_i)}$$

$i$: Species identity
$c_i$: Parts of abundance common to both sites
$u_i$: Parts of abundance unique to either sites

**(b)** Types of losses and gains in species abundance

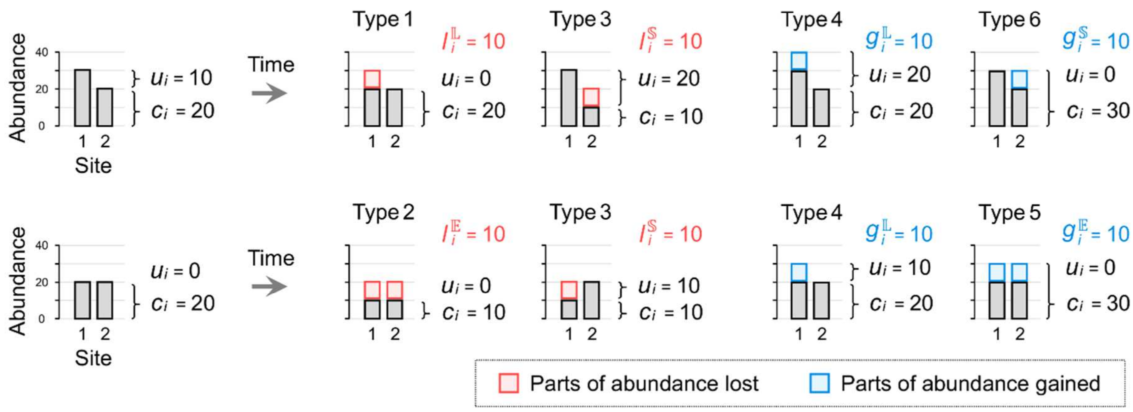


**(c)** Consequences of abundance losses and gains on dissimilarity

| Type | Description | Temporal changes in … $c_i$ | $u_i$ | $\beta_{\text{Ruž}}$ and $\beta_{\text{BC}}$ |
|---|---|---|---|---|
| 1 | Abundance ($a_i$) decreases in the site where $a_i$ is larger ($\mathbb{L}$) than that in the other site. | 0 | $-l_i^{\mathbb{L}}$ | − |
| 2 | $a_i$ decreases by equal amount ($\mathbb{E}$) in both sites. | $-l_i^{\mathbb{E}}$ | 0 | + |
| 3 | $a_i$ decreases in the site where $a_i$ is smaller ($\mathbb{S}$) than, or equal to, that in the other site. | $-l_i^{\mathbb{S}}$ | $+l_i^{\mathbb{S}}$ | + |
| 4 | $a_i$ increases in the site where $a_i$ is larger ($\mathbb{L}$) than, or equal to, that in the other site. | 0 | $+g_i^{\mathbb{L}}$ | + |
| 5 | $a_i$ increases by equal amount ($\mathbb{E}$) in both sites. | $+g_i^{\mathbb{E}}$ | 0 | − |
| 6 | $a_i$ increases in the site where $a_i$ is smaller ($\mathbb{S}$) than that in the other site. | $+g_i^{\mathbb{S}}$ | $-g_i^{\mathbb{S}}$ | − |

**Figure S1.** Schematic representation of the temporal changes in spatial beta diversity and types of losses and gains in species abundances that drive the changes. (a) Ružička and Bray–Curtis dissimilarity indices. (b) Examples of abundances losses and gains categorized into six types based on their impacts on beta diversity. (c) Descriptions of the six types of abundances losses and gains.

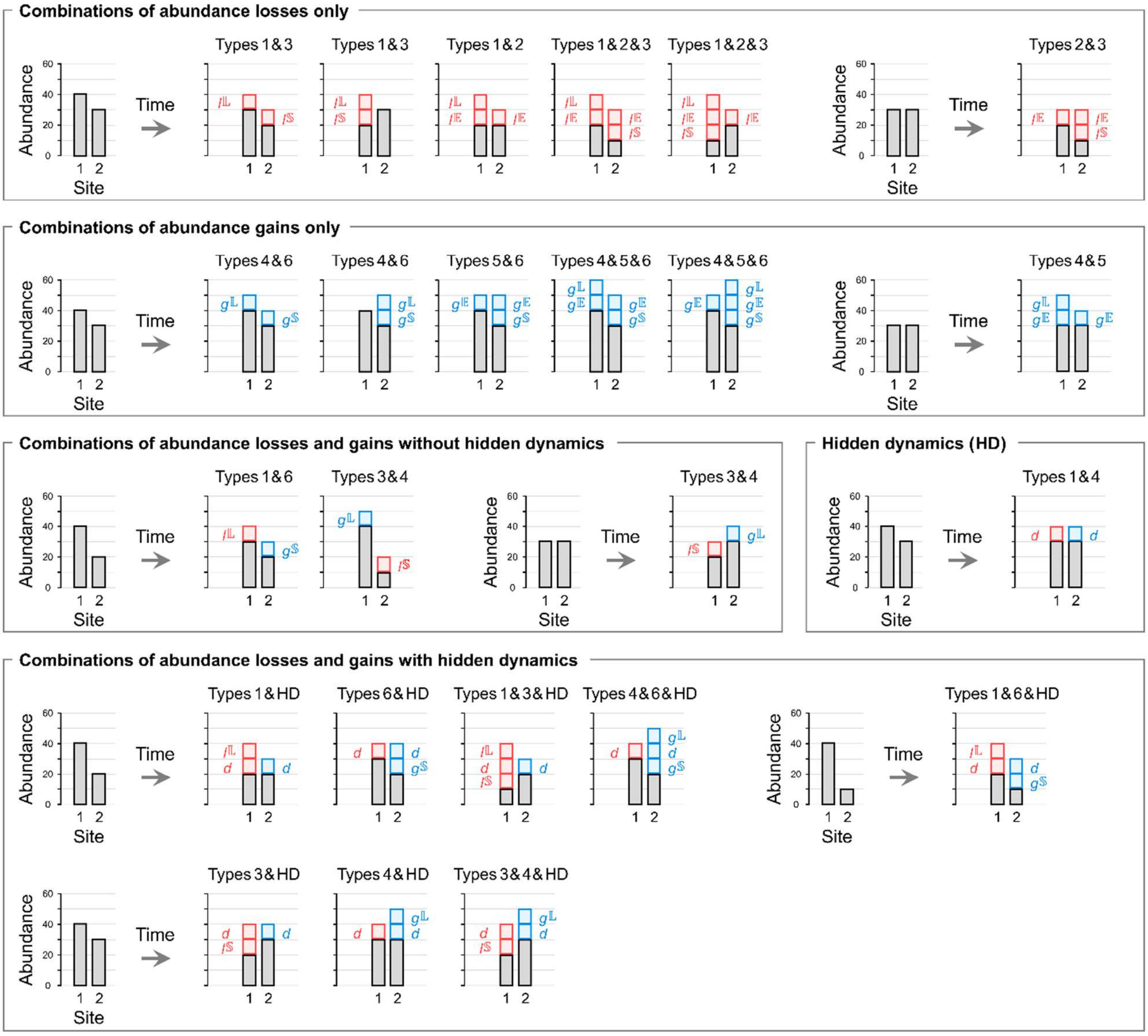


**Figure S2.** Possible combinations of abundance losses and gains including hidden dynamics in the case of two sites. In total, there are 32 possible ways of abundance losses and gains; i.e., the 8 types shown in Figure S1 and the 24 combinations shown here.

**Appendix S2: Sensitivity analyses**

We conducted sensitivity analyses to verify the impacts of losses and gains in species abundances on $\Delta\beta$ components ($\Delta\beta_1$, $\Delta\beta_2$, $\Delta\beta_3$, $\Delta\beta_4$, $\Delta\beta_5$, $\Delta\beta_6$, $\Delta\beta_{L-}$, $\Delta\beta_{L+}$, $\Delta\beta_{G-}$, and $\Delta\beta_{G+}$). We varied nine variables $L^{\mathbb{L}}$, $L^{\mathbb{E}}$, $L^{\mathbb{S}}$, $G^{\mathbb{L}}$, $G^{\mathbb{E}}$, $G^{\mathbb{S}}$, $D$, $C$, and $U$ one by one from 0 to 100 while fixing the others at 5. We then calculated the sizes of the components based on Ružička and Bray–Curtis indices ($\beta_{Ruž}$ and $\beta_{BC}$). Note that the components of $\beta_{BC}$ and the normalized abundance-based Whittaker's beta ($\beta_W$) are mathematically identical (see Appendix S3). The sensitivity analyses we conducted here for abundance losses and gains are analogous to what we did for species extinctions and colonizations using presence–absence data (Tatsumi et al., 2021).

The sensitivity analyses confirmed that the component $\Delta\beta_{L-}$ ($\Delta\beta_1$) decreases with $L^{\mathbb{L}}$ (the loss in species abundance unique to either site), whereas the component $\Delta\beta_{L+}$ ($\Delta\beta_2$ and $\Delta\beta_3$) increases with $L^{\mathbb{E}}$ and $L^{\mathbb{S}}$ (the loss in species abundance common to both sites) (Figures S3a–c, S4a–c). Similarly, for colonization, $\Delta\beta_{G+}$ ($\Delta\beta_4$) increases with $G^{\mathbb{L}}$ (the gain in species abundance unique to either site), whereas $\Delta\beta_{G-}$ ($\Delta\beta_5$ and $\Delta\beta_6$) decreases with $G^{\mathbb{E}}$ and $G^{\mathbb{S}}$ (the gain in species abundance common to both sites) (Figures S3d–f, S4d–f). These results were similar to what we found with the portioning of extinction and colonization components (Tatsumi et al., 2021).

We also found that $\Delta\beta_3$ and $\Delta\beta_6$ change more largely than $\Delta\beta_1$, $\Delta\beta_2$, $\Delta\beta_4$, and $\Delta\beta_5$ do in response to the varying values of $L^{\mathbb{L}}$, $L^{\mathbb{E}}$, $L^{\mathbb{S}}$, $G^{\mathbb{L}}$, $G^{\mathbb{E}}$, $G^{\mathbb{S}}$ (Figures S3a–f, S4a–f). This finding agrees well with the fact that $\Delta\beta_3$ and $\Delta\beta_6$ reflect numerical changes in both unique and shared parts of species abundances (see types 3 and 6 in Figure S1), whereas the other $\Delta\beta$ components are only responsible for either unique or shared species (types 1, 2, 4, and 5). Furthermore, consistent with its definition, the hidden species dynamics ($D$) simultaneously decreased $\Delta\beta_1$ and increased $\Delta\beta_3$ without affecting the total $\Delta\beta$ (Figures S3g, S4g). The amount of species abundance common to both sites (constant $C$) caused no impact on $\Delta\beta$ but reduced the relative importance of other species (Figures S3h, S4h). Similarly, the amount of species abundance unique to either site (constant $U$) caused no impact on $\Delta\beta$ (Figures S3i, S4i). Again, these results were similar to what we found with the portioning of extinction and colonization components (Tatsumi et al., 2021).

**Reference**

Tatsumi, S., Iritani, R., Cadotte, M.W. (2021). Temporal changes in spatial variation: partitioning the extinction and colonisation components of beta diversity. *Ecology Letters* 24, 1063–1072.

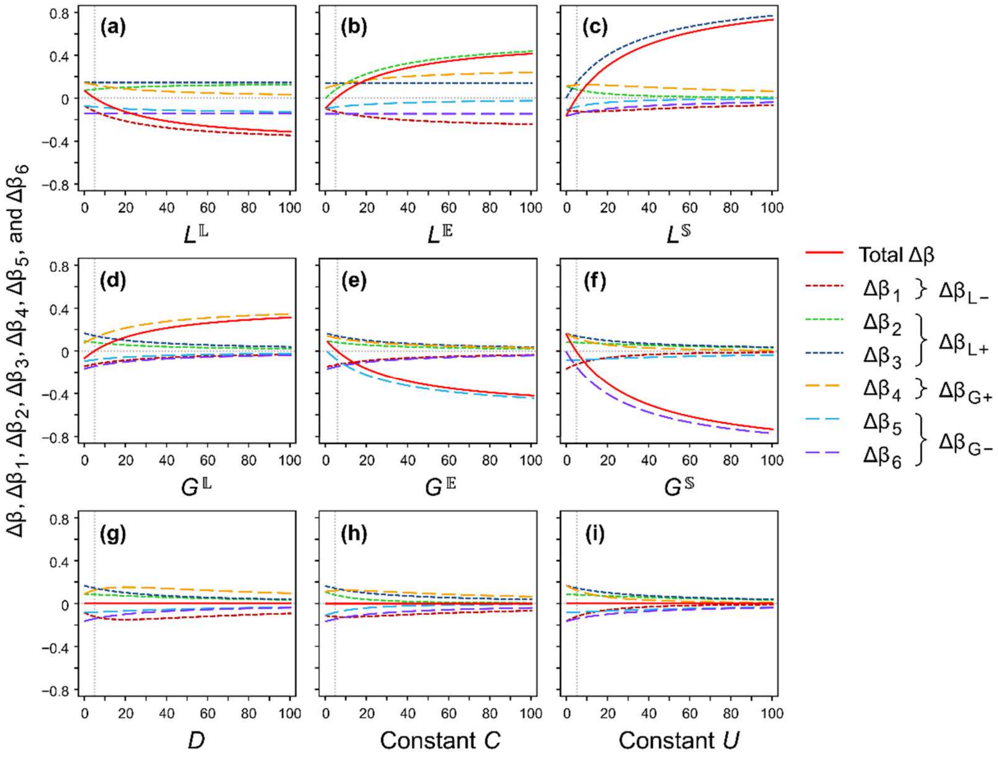


**Figure S3.** Sensitivity analyses of the response of temporal changes in Ružička index and its components to varying amount of species abundance common to either site and unique to both sites.

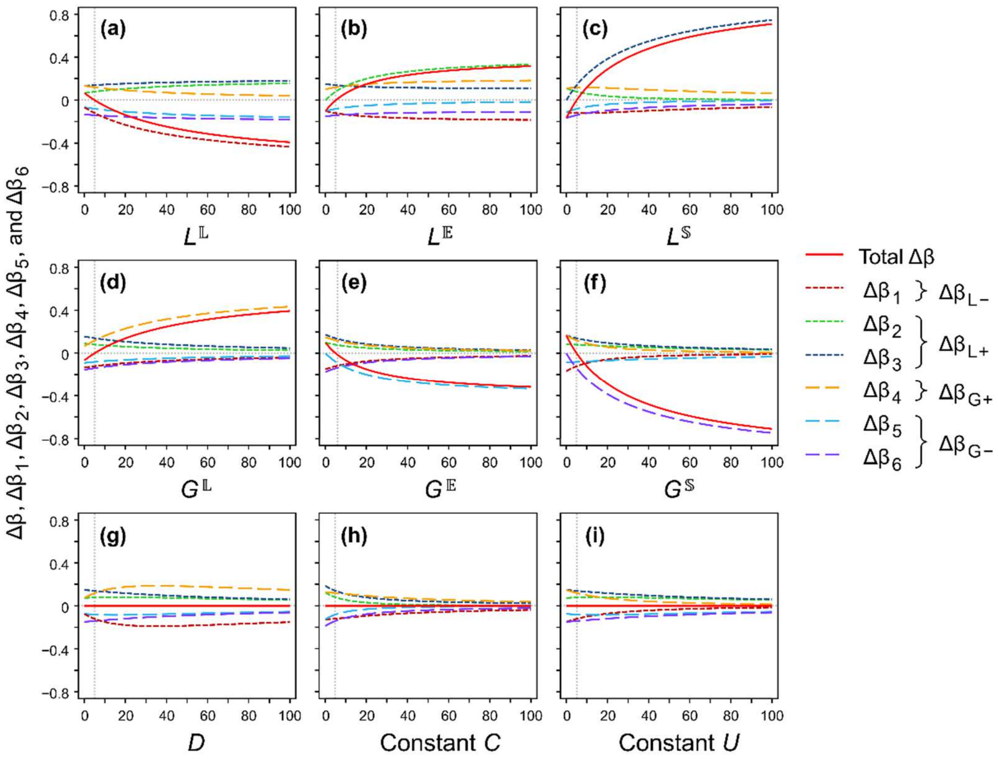


**Figure S4.** Sensitivity analyses of the response of temporal changes in Bray–Curtis index (which is mathematically identical to the normalized abundance-based Whittaker's beta when $N = 2$) and its components to varying amount of species abundance common to either site and unique to both sites.

**Appendix S3: Derivation of partitioning equations: Multisite variation**

Our partitioning approach is applicable to multisite measures as well. For cases with more than two sites, averaging pairwise dissimilarities ($\beta_{Ruž}$ and $\beta_{BC}$) across all pairs of sites is a suboptimal approach due to the lack of their statistical independence (Baselga, 2010, 2017).

Here, we consider the normalized abundance-based Whittaker's beta ($\beta_W$) proposed by Baselga (2017). Harrison et al. (1992) defined the normalized incidence-based Whittaker's beta as $(\gamma/\bar{\alpha} - 1)/(N - 1)$, where $\gamma$ is the regional species richness, $\bar{\alpha}$ is the mean local species richness, and $N$ is the number of sites. While the original Whittaker's beta ($\gamma/\bar{\alpha}$) ranges in the interval [1, $N$] (Whittaker, 1960), the normalized measure ranges in the interval [0, 1] (Harrison et al., 1992). This normalized measure has been extended by Baselga (2017) to account for abundance. $\beta_W$ and other variables are defined as follows:

| | |
|---|---|
| $\beta_W = \frac{N \cdot \sum_i^S \max_k\{a_{ik}\} - A}{(N-1)A}$ | Baselga's beta, or the normalized abundance-based Whittaker's beta. |
| $N$ | Number of sites. |
| $S$ | Number of species. |
| $a_{ik}$ | Abundance of species $i$ in site $k$. |
| $\max_k\{a_{ik}\}$ | The maximum abundance of species $i$ across all the sites. Also denoted as $m_i$. |
| $A$ | Total abundance of all species in all sites (i.e., $A = \sum_{i=1}^S \sum_{k=1}^N a_{ik}$). |
| $a_{i+}$ | Abundance of species $i$ summed across all sites (i.e., $a_{i+} = \sum_{k=1}^N a_{ik}$). |
| $l_{in}^{\mathbb{L}}$ | Amount of loss in the maximum abundance of species $i$ ($m_i$) in $n$ sites. |
| $l_{in}^{\mathbb{S}}$ | Amount of loss in $a_{i+}$ without causing any change in $m_i$ — i.e., abundance loss that does not affect the maximum abundance of species $i$ in $n$ sites. |
| $g_{in}^{\mathbb{L}}$ | Amount of gain in the maximum abundance of species $i$ ($m_i$) in $n$ sites. |
| $g_{in}^{\mathbb{S}}$ | Amount of gain in $a_{i+}$ without causing any change in $m_i$ — i.e., abundance gain that does not affect the maximum abundance of species $i$ in $n$ sites. |
| $d_{ivw}$ | Amount of losses in the abundance of species $i$ in $v$-number of sites and the offsetting gains in $w$-number of sites occurring within the same time interval. |

We extend the variables $l_i^{\mathbb{L}}$, $l_i^{\mathbb{E}}$, $l_i^{\mathbb{S}}$, $g_i^{\mathbb{L}}$, $g_i^{\mathbb{E}}$, and $g_i^{\mathbb{S}}$, defined earlier (Appendix S1; Figures S1, S2) to denote the temporal changes in the maximum abundance ($m_i$) and the total abundance ($a_{i+}$) of species $i$. First, we extend $l_i^{\mathbb{L}}$ to denote the decrease in species abundance associated with both $m_i$ and $a_{i+}$. This type of abundance loss decreases $m_i$ by $l_i^{\mathbb{L}}$. For example, $m_i$ decreases by 10 when $l_i^{\mathbb{L}} = 10$. The amount of decrease in $a_{i+}$, on the other hand, depends on the number of sites in which a given species $i$ showed its maximum abundance at $t = 1$. This dependency can be taken explicitly into account by adding a subscript $n$ to $l_i^{\mathbb{L}}$, such that $l_{in}^{\mathbb{L}}$ denotes the decrease in the maximum abundance across $n$ sites. This type of abundance loss decreases $a_{i+}$ by $n \cdot l_{in}^{\mathbb{L}}$. For instance, $a_{i+}$ decreases by 10 when $l_{i1}^{\mathbb{L}} = 10$ and decreases by 20 when $l_{i2}^{\mathbb{L}} = 10$, whereas $m_i$ decreases by 10 in either case. Similarly, we define $g_{in}^{\mathbb{L}}$ to denote the increase in the maximum abundance of species $i$ across $n$ sites.

We extend the definition of $l_i^{\mathbb{E}}$ to denote the abundance loss in all sites ($N$) across which a given species $i$ showed its maximum abundance at $t = 1$. In this case, $m_i$ and $a_{i+}$ decrease by $l_i^{\mathbb{E}}$ and $N \cdot l_i^{\mathbb{E}}$, respectively. Thus, $l_i^{\mathbb{E}}$ can be seen as a special case of $l_{in}^{\mathbb{L}}$ when $n = N$. We therefore rewrite $l_i^{\mathbb{E}}$ as $l_{iN}^{\mathbb{L}}$ hereafter. Similarly, we rewrite $g_i^{\mathbb{E}}$ as $g_{iN}^{\mathbb{L}}$ for the gain in abundance across all sites in which a given species $i$ showed its maximum abundance at $t = 2$.

We define $l_i^{\mathbb{S}}$ as the abundance loss associated with the decrease in $a_{i+}$ but not $m_i$. We use the subscript $n$ to let $l_{in}^{\mathbb{S}}$ denote the abundance loss in $n$ sites across which a given species $i$ showed smaller abundance than, or equal abundance to, $m_i$ at both $t = 1$ and 2. This type of abundance loss decreases $a_{i+}$ by $n \cdot l_{in}^{\mathbb{S}}$ but does not alter $m_i$. Similarly, we define $g_{in}^{\mathbb{S}}$ as the increase in abundance across $n$ sites where species $i$ showed smaller abundance than, or equal abundance to, $m_i$ at both $t = 1$ and 2.

Hidden dynamics can occur not only in the case of two sites ($N = 2$) but also multiple sites ($N \geq 3$). To account for this, we extend the definition of hidden dynamics as the abundance losses in the site(s) where a given species $i$ showed its maximum abundance at $t = 1$ and the offsetting gains in the other site(s). By adding subscripts $v$ and $w$ to $d_i$, we write $d_{ivw}$ for the abundance losses in $v$-number of sites and the offsetting gains in $w$-number of sites occurring within the same time interval ($v \geq 1$, $w \geq 1$, and $v + w \leq N$).

Again, for brevity, we write the sum of a given variable across all species (1, 2, ..., $S$) using the uppercase letters (e.g., $\sum_{i=1}^{S} \max_k\{a_{ik}\} = \sum_{i=1}^{S} m_i = M$, $\sum_{i=1}^{S} l_{in}^{\mathbb{L}} = L_n^{\mathbb{L}}$, and $\sum_{i=1}^{S} d_{ivw} = D_{vw}$). Equation 1 in the main text can be rewritten using the extended definitions by replacing $L^{\mathbb{L}}$, $L^{\mathbb{E}}$, $L^{\mathbb{S}}$, $G^{\mathbb{L}}$, $G^{\mathbb{E}}$, $G^{\mathbb{S}}$, and $D$ with $L_1^{\mathbb{L}}$, $L_2^{\mathbb{L}}$, $L_1^{\mathbb{S}}$, $G_1^{\mathbb{L}}$, $G_2^{\mathbb{L}}$, $G_1^{\mathbb{S}}$, and $D_{11}$, respectively.

#### *Two-site cases*

To partition $\beta_W$, we first consider the case of two communities ($N = 2$). The temporal changes in $M$ and $A$ can be written as

$$\Delta M = M' - M = -L_1^{\mathbb{L}} - L_2^{\mathbb{L}} + G_1^{\mathbb{L}} + G_2^{\mathbb{L}}$$
$$\Delta A = A' - M = -L_1^{\mathbb{L}} - 2L_2^{\mathbb{L}} - L_1^{\mathbb{S}} + G_1^{\mathbb{L}} + 2G_2^{\mathbb{L}} + G_1^{\mathbb{S}}.$$

The temporal changes in $\beta_W$ when $N = 2$ can then be partitioned as

$$\Delta\beta_W = \beta'_W - \beta_W$$
$$= \frac{NM' - A'}{(N-1)A'} - \frac{NM - A}{(N-1)A}$$

$$
\begin{aligned}
&= \frac{NM'A - NMA'}{(N-1)AA'} \\
&= \frac{NM\left(\frac{\Delta M}{M} - \frac{\Delta A}{A}\right)}{(N-1)A'} \qquad \because \; M' = M + \Delta M, A' = A + \Delta A \\
&= \underbrace{\frac{NM}{(N-1)A'}}_{q} \times \left( \frac{-L_1^{\mathbb{L}} - L_2^{\mathbb{L}} + G_1^{\mathbb{L}} + G_2^{\mathbb{L}}}{M} + \frac{L_1^{\mathbb{L}} + 2L_2^{\mathbb{L}} + L_1^{\mathbb{S}} - G_1^{\mathbb{L}} - 2G_2^{\mathbb{L}} - G_1^{\mathbb{S}}}{A} \right),
\end{aligned}
$$

where $M$ and $A$ are both non-zero. Accordingly, we get the partitioning equation, without the hidden dynamics, which reads

$$
\begin{aligned}
\Delta\beta_{\mathrm{W}} &= \beta'_{\mathrm{W}} - \beta_{\mathrm{W}} \\
&= \underbrace{\left(\frac{-q}{M} + \frac{q}{A}\right)L_1^{\mathbb{L}}}_{\Delta\beta_1} + \underbrace{\left(\frac{-q}{M} + \frac{2q}{A}\right)L_2^{\mathbb{L}}}_{\Delta\beta_2} + \underbrace{\frac{q}{A}L_1^{\mathbb{S}}}_{\Delta\beta_3} + \underbrace{\left(\frac{q}{M} - \frac{q}{A}\right)G_1^{\mathbb{L}}}_{\Delta\beta_4} + \underbrace{\left(\frac{q}{M} - \frac{2q}{A}\right)G_2^{\mathbb{L}}}_{\Delta\beta_5} + \underbrace{\frac{-q}{A}G_1^{\mathbb{S}}}_{\Delta\beta_6}.
\end{aligned}
$$

As we did for pairwise dissimilarity ($\beta_{\mathrm{Ruž}}$ and $\beta_{\mathrm{BC}}$), the hidden dynamics can be accounted for by adding $D$ to the multipliers of the first and fourth terms:

$$
\begin{aligned}
\Delta\beta_{\mathrm{W}} &= \beta'_{\mathrm{W}} - \beta_{\mathrm{W}} \\
&= \underbrace{\left(\frac{-q}{M} + \frac{q}{A}\right)\left(L_1^{\mathbb{L}} + D_{11}\right)}_{\Delta\beta_1} + \underbrace{\left(\frac{-q}{M} + \frac{2q}{A}\right)L_2^{\mathbb{L}}}_{\Delta\beta_2} + \underbrace{\frac{q}{A}L_1^{\mathbb{S}}}_{\Delta\beta_3} + \underbrace{\left(\frac{q}{M} - \frac{q}{A}\right)\left(G_1^{\mathbb{L}} + D_{11}\right)}_{\Delta\beta_4} + \underbrace{\left(\frac{q}{M} - \frac{2q}{A}\right)G_2^{\mathbb{L}}}_{\Delta\beta_5} + \underbrace{\frac{-q}{A}G_1^{\mathbb{S}}}_{\Delta\beta_6}.
\end{aligned}
$$

We define the six terms as the loss and gain components of temporal changes in the normalized abundance-based Whittaker's beta ($\Delta\beta_{\mathrm{W}}$) at $N = 2$.

Note that $\beta_{\mathrm{W}}$ and $\beta_{\mathrm{BC}}$ are identical when $N = 2$ (Beselga, 2017):

$$
\begin{aligned}
\beta_{\mathrm{W}} &= \frac{NM - A}{(N-1)A} \\
&= \frac{2(C+U) - (2C+U)}{2C+U} \qquad \because \; M = C + U, A = 2C + U \\
&= \frac{U}{2C+U} = \beta_{\mathrm{BC}}.
\end{aligned}
$$

The six components of $\Delta\beta_{\mathrm{W}}$ and $\Delta\beta_{\mathrm{BC}}$ of ($\Delta\beta_1$, $\Delta\beta_2$, $\Delta\beta_3$, $\Delta\beta_4$, $\Delta\beta_5$, and $\Delta\beta_6$) are also identical to each other when $N = 2$:

$$
\begin{aligned}
\Delta\beta_1 &= \left(\frac{-q}{M} + \frac{q}{A}\right)L_1^{\mathbb{L}} \\
&= \left(\frac{-2}{A'} + \frac{2M}{AA'}\right)L_1^{\mathbb{L}} = \frac{2M - 2A}{AA'}L_1^{\mathbb{L}} = \frac{-2C}{(2C+U)(2C'+U')}L_1^{\mathbb{L}} \\
&= -\frac{p}{U}L^{\mathbb{L}}
\end{aligned}
$$

$$
\Delta\beta_2 = \left(\frac{-q}{M} + \frac{2q}{A}\right)L_2^{\mathbb{L}}
$$

$$= \left(\frac{-2}{A'} + \frac{4M}{AA'}\right) L_2^{\mathbb{L}} = \frac{2U}{(2C+U)(2C'+U')} L_2^{\mathbb{L}}$$
$$= \frac{p}{C} L^{\mathbb{E}}$$

$$\Delta\beta_3 = \frac{q}{A} L_1^{\mathbb{S}}$$
$$= \frac{2M}{AA'} L_1^{\mathbb{S}} = \frac{2C+2U}{(2C+U)(2C'+U')} L_1^{\mathbb{S}}$$
$$= \left(\frac{p}{U} + \frac{p}{C}\right) L^{\mathbb{S}}.$$

We also obtain $\Delta\beta_4 = \left(\frac{q}{M} - \frac{q}{A}\right) G_1^{\mathbb{L}} = \frac{p}{U} G^{\mathbb{L}}$, $\Delta\beta_5 = \left(\frac{q}{M} - \frac{q}{A}\right) G_1^{\mathbb{L}} = \frac{p}{U} G^{\mathbb{L}}$, and $\Delta\beta_6 = \left(\frac{q}{M} - \frac{q}{A}\right) G_1^{\mathbb{L}} = \frac{p}{U} G^{\mathbb{L}}$ by replacing $L_1^{\mathbb{L}}$ with $-G_1^{\mathbb{L}}$, $L_2^{\mathbb{L}}$ with $-G_2^{\mathbb{L}}$, and $L_1^{\mathbb{S}}$ with $-G_1^{\mathbb{S}}$ in the above equations, respectively.

***Three-site cases***

Let us next expand the $\beta_W$-based partitioning equations to the case of three sites. For the sake of simplicity, we first leave out the hidden dynamics. When $N = 3$, there will be one, two or three site(s) in which species $i$ shows its maximum abundance ($m_i$). The subscript $n$ of $l_{in}^{\mathbb{L}}$ and $l_{in}^{\mathbb{L}}$ will thus range from 1 to 3: $l_{i1}^{\mathbb{L}}$, $l_{i2}^{\mathbb{L}}$, $l_{i3}^{\mathbb{L}}$, $g_{i1}^{\mathbb{L}}$, $g_{i2}^{\mathbb{L}}$, and $g_{i3}^{\mathbb{L}}$. On the other hand, there will be one or two site(s) in which species $i$ shows abundance less than $m_i$. The temporal changes in species abundance that do not affect $m_i$ can thus occur in 1 or 2 sites: $l_{i1}^{\mathbb{S}}$, $l_{i2}^{\mathbb{S}}$, $g_{i1}^{\mathbb{S}}$, and $g_{i2}^{\mathbb{S}}$. As such, we get

$$\Delta M = M' - M = -L_1^{\mathbb{L}} - L_2^{\mathbb{L}} - L_3^{\mathbb{L}} + G_1^{\mathbb{L}} + G_2^{\mathbb{L}} + G_3^{\mathbb{L}}$$
$$\Delta A = A' - A = -L_1^{\mathbb{L}} - 2L_2^{\mathbb{L}} - 3L_3^{\mathbb{L}} - L_1^{\mathbb{S}} - 2L_2^{\mathbb{S}} + G_1^{\mathbb{L}} + 2G_2^{\mathbb{L}} + 3G_3^{\mathbb{L}} + G_1^{\mathbb{S}} + 2G_2^{\mathbb{S}}.$$

The temporal changes in $\beta_W$ when $N = 3$ can thus be partitioned as

$$\Delta\beta_W = \beta'_W - \beta_W$$
$$= \underbrace{\frac{NM}{(N-1)A'}}_{q} \times \left(\frac{\Delta M}{M} - \frac{\Delta A}{A}\right)$$
$$= \underbrace{\left(\frac{-q}{M} + \frac{q}{A}\right) L_1^{\mathbb{L}}}_{\Delta\beta_{L-}} + \underbrace{\left(\frac{-q}{M} + \frac{2q}{A}\right) L_2^{\mathbb{L}}}_{\Delta\beta_{L-} \text{ or } \Delta\beta_{L+}} + \underbrace{\left(\frac{-q}{M} + \frac{3q}{A}\right) L_3^{\mathbb{L}} + \frac{q}{A} L_1^{\mathbb{S}} + \frac{2q}{A} L_2^{\mathbb{S}}}_{\Delta\beta_{L+}}$$
$$+ \underbrace{\left(\frac{q}{M} - \frac{q}{A}\right) G_1^{\mathbb{L}}}_{\Delta\beta_{G+}} + \underbrace{\left(\frac{q}{M} - \frac{2q}{A}\right) G_2^{\mathbb{L}}}_{\Delta\beta_{G-} \text{ or } \Delta\beta_{G+}} + \underbrace{\left(\frac{q}{M} - \frac{3q}{A}\right) G_3^{\mathbb{L}} + \frac{-q}{A} G_1^{\mathbb{S}} + \frac{-2q}{A} G_2^{\mathbb{S}}}_{\Delta\beta_{G-}}$$

Here, the terms are no longer categorized *a priori* into $\Delta\beta_1$, $\Delta\beta_2$, $\Delta\beta_3$, $\Delta\beta_4$, $\Delta\beta_5$, and $\Delta\beta_6$. This is because when $N \geq 3$, some of the terms take either negative or positive values depending on the sizes of $A$ and $M$. The terms can, however, be summed by groups in such a way that they represent homogenization and differentiation driven by abundance losses and gains ($\Delta\beta_{L-}$, $\Delta\beta_{L+}$, $\Delta\beta_{G-}$, and $\Delta\beta_{G+}$).

To explain why some terms can be either negative or positive, let us take the terms $\left(\frac{-q}{M} + \frac{q}{A}\right) L_1^{\mathbb{L}}$,

$\left(\frac{-q}{M}+\frac{2q}{A}\right)L_2^{\mathbb{L}}$, and $\left(\frac{-q}{M}+\frac{3q}{A}\right)L_3^{\mathbb{L}}$ as examples. The variables $L_1^{\mathbb{L}}$, $L_2^{\mathbb{L}}$, and $L_3^{\mathbb{L}}$ indicate the sum of decreases in $m_i$ in one, two, and three sites, respectively. The relative size of $A$ to $M$ is minimized when each species is present only in one site ($A = M$) and is maximized when each species shows the same abundance across all three sites (i.e., maximum abundances found in all sites; $A = 3M$). Thus, $\left(\frac{-q}{M}+\frac{q}{A}\right)L_1^{\mathbb{L}}$ is always nonpositive and $\left(\frac{-q}{M}+\frac{3q}{A}\right)L_3^{\mathbb{L}}$ is always nonnegative. These mathematical results agree well with our intuition that the loss of species abundance unique to a single site results in biotic homogenization, whereas the loss of species abundance common to all sites leads to differentiation (Socolar et al., 2016; Tatsumi et al., 2020, 2021). On the other hand, $\left(\frac{-q}{M}+\frac{2q}{A}\right)L_2^{\mathbb{L}}$ can change its sign depending on the balance between $A$ and $M$. This reflects the fact that the loss of species abundance unique to an intermediate number of sites can either drive homogenization or differentiation depending on the abundance distribution of all species across all sites.

We now consider the hidden dynamics when $N = 3$. As described above, when $N = 2$, there is only one possible combination of abundance loss and gain that could occur within the same time interval — i.e., an abundance loss in the site where species $i$ showed its maximum abundance ($m_i$) and an offsetting abundance gain in the other site. When $N = 3$, the number of possible combinations increases to three — i.e., an abundance loss(es) in one or two site(s) where species $i$ showed $m_i$ and an offsetting abundance gain(s) in the other one or two site(s). We denote the number of sites that corresponds such abundance losses and gains as $v$ and $w$, respectively, where $v \geq 1$, $w \geq 1$, and $v + w \leq N$.

We write $D_{vw}$ for the abundance losses in $v$-number of sites and the offsetting gains in $w$-number of sites occurring within the same time interval. The amount of change in $\Delta\beta$ driven by $D_{vw}$ is equivalent to that cause by $L_v^{\mathbb{L}}$ and $G_w^{\mathbb{L}}$. By adding the three possible $D_{pq}$ (i.e., $D_{11}$, $D_{12}$, and $D_{21}$) to the corresponding terms in the equation above, the temporal changes in $\beta_{\mathrm{W}}$ when $N = 3$ can be partitioned as

$$\begin{aligned}\Delta\beta_{\mathrm{W}} &= \beta'_{\mathrm{W}} - \beta_{\mathrm{W}} \\ &= \underbrace{\left(\frac{-q}{M}+\frac{q}{A}\right)\left(L_1^{\mathbb{L}} + D_{11} + D_{12}\right)}_{\Delta\beta_{\mathrm{L-}}} + \underbrace{\left(\frac{-q}{M}+\frac{2q}{A}\right)\left(L_2^{\mathbb{L}} + D_{21}\right)}_{\Delta\beta_{\mathrm{L-}} \text{ or } \Delta\beta_{\mathrm{L+}}} + \underbrace{\left(\frac{-q}{M}+\frac{3q}{A}\right)L_3^{\mathbb{L}} + \frac{q}{A}L_1^{\mathbb{S}} + \frac{2q}{A}L_2^{\mathbb{S}}}_{\Delta\beta_{\mathrm{L+}}} \\ &\quad + \underbrace{\left(\frac{q}{M}-\frac{q}{A}\right)\left(G_1^{\mathbb{L}} + D_{11} + D_{21}\right)}_{\Delta\beta_{\mathrm{G+}}} + \underbrace{\left(\frac{q}{M}-\frac{2q}{A}\right)\left(G_2^{\mathbb{L}} + hD_{12}\right)}_{\Delta\beta_{\mathrm{G-}} \text{ or } \Delta\beta_{\mathrm{G+}}} + \underbrace{\left(\frac{q}{M}-\frac{3q}{A}\right)G_3^{\mathbb{L}} + \frac{-q}{A}G_1^{\mathbb{S}} + \frac{-2q}{A}G_2^{\mathbb{S}}}_{\Delta\beta_{\mathrm{G-}}}.\end{aligned}$$

#### *Generalization: N-site cases*

We now generalize the $\beta_{\mathrm{W}}$-based partitioning equations to the case of $N$ sites. Again, for the sake of simplicity, we first leave out the hidden dynamics. When $N$, there will be 1, 2, ..., $N$ sites in which species $i$ shows its maximum abundance ($m_i$). Thus, as we saw in the case of $N = 3$, the subscript $n$ of $l_{in}^{\mathbb{L}}$ and $l_{in}^{\mathbb{L}}$ will range from 1 to $N$: $l_{i1}^{\mathbb{L}}$, $l_{i2}^{\mathbb{L}}$, ..., $l_{in}^{\mathbb{L}}$ and $g_{i1}^{\mathbb{L}}$, $g_{i2}^{\mathbb{L}}$, ..., $g_{in}^{\mathbb{L}}$. On the other hand, there will be 1, 2, ..., $N - 1$ sites in which species $i$ shows abundance less than $m_i$. The temporal changes in species abundance that do not affect $m_i$ can thus occur in 1, 2, ..., $N-1$ sites: $l^{\mathbb{S}}_{i1}$, $l^{\mathbb{S}}_{i2}$, ..., $l^{\mathbb{S}}_{i(N-1)}$ and $g^{\mathbb{S}}_{i1}$, $g^{\mathbb{S}}_{i2}$, ..., $g^{\mathbb{S}}_{i(N-1)}$. We thus get

$$\Delta M = M' - M = -\sum_{n=1}^{N} L^{\mathbb{L}}_n + \sum_{n=1}^{N} G^{\mathbb{L}}_n$$

$$\Delta A = A' - A = -\sum_{n=1}^{N} nL^{\mathbb{L}}_n - \sum_{n=1}^{N-1} nL^{\mathbb{S}}_n + \sum_{n=1}^{N} nG^{\mathbb{L}}_n + \sum_{n=1}^{N-1} nG^{\mathbb{S}}_n.$$

The temporal changes in $\beta_W$ can thus be partitioned as

$$\begin{aligned}\Delta\beta_W &= \beta'_W - \beta_W \\ &= \underbrace{\frac{NM}{(N-1)A'}}_{q} \times \left(\frac{\Delta M}{M} - \frac{\Delta A}{A}\right) \\ &= \underbrace{\sum_{n=1}^{N}\left(\frac{-q}{M} + \frac{nq}{A}\right)L^{\mathbb{L}}_n}_{\Delta\beta_{L-}\text{ or }\Delta\beta_{L+}} + \underbrace{\sum_{n=1}^{N-1}\frac{nq}{A}L^{\mathbb{S}}_n}_{\Delta\beta_{L+}} + \underbrace{\sum_{n=1}^{N}\left(\frac{q}{M} - \frac{nq}{A}\right)L^{\mathbb{L}}_n}_{\Delta\beta_{G-}\text{ or }\Delta\beta_{G+}} + \underbrace{\sum_{n=1}^{N-1}\frac{-nq}{A}L^{\mathbb{S}}_n}_{\Delta\beta_{G-}}.\end{aligned}$$

Finally, let us consider the generalization of the hidden dynamics. The hidden dynamics can occur in such a way that the abundance of species $i$ decreases in 1, 2, ..., or $N-1$ sites where it showed is maximum abundance ($m_i$) and increases in the other 1, 2, ..., or $N-1$ sites. As defined above, we write $D_{vw}$ for the abundance losses in $v$-number of sites and the offsetting gains in $w$-number of sites occurring within the same time interval. The possible combinations of $v$ and $w$ that meet the conditions $v \geq 1$, $w \geq 1$, and $v + w \leq N$ equals $\sum_{x=1}^{1} x + \sum_{x=1}^{2} x + \sum_{x=1}^{3} x + \cdots + \sum_{x=1}^{(N-1)} x$. By adding all the possible $D_{vw}$ (i.e., $D_{11}$, $D_{12}$, $D_{13}$, ..., $D_{1(N-1)}$, $D_{21}$, $D_{22}$, ..., $D_{2(N-2)}$, $D_{31}$, ..., $D_{(N-2)1}$, $D_{(N-2)2}$, $D_{(N-1)1}$) to the corresponding multipliers ($L^{\mathbb{L}}_v$ and $G^{\mathbb{L}}_w$) in the equation above, the temporal changes in $\beta_W$ at any number of sites ($N$) can be partitioned as

$$\begin{aligned}\Delta\beta_W &= \beta'_W - \beta_W \\ &= \underbrace{\sum_{n=1}^{N}\left(\frac{-q}{M} + \frac{nq}{A}\right)L^{\mathbb{L}}_n + \sum_{\{v,w|v+w\leq N\}}\left(\frac{-q}{M} + \frac{vq}{A}\right)D_{vw}}_{\Delta\beta_{L-}\text{ or }\Delta\beta_{L+}} + \underbrace{\sum_{n=1}^{N-1}\frac{nq}{A}L^{\mathbb{S}}_n}_{\Delta\beta_{L+}} \\ &\quad + \underbrace{\sum_{n=1}^{N}\left(\frac{q}{M} - \frac{nq}{A}\right)G^{\mathbb{L}}_n + \sum_{\{v,w|v+w\leq N\}}\left(\frac{q}{M} - \frac{wq}{A}\right)D_{vw}}_{\Delta\beta_{G-}\text{ or }\Delta\beta_{G+}} + \underbrace{\sum_{n=1}^{N-1}\frac{-nq}{A}G^{\mathbb{S}}_n}_{\Delta\beta_{G-}}\end{aligned}$$

The terms $\Delta\beta_{L-}$ and $\Delta\beta_{L+}$ indicate homogenization and differentiation, respectively, caused by abundance losses. Similarly, $\Delta\beta_{G-}$ and $\Delta\beta_{G+}$ represent homogenization and differentiation, respectively, caused by abundance gains.

### References

Baselga, A. (2010). Partitioning the turnover and nestedness components of beta diversity. *Global*

*Ecology and Biogeography*, *19*, 134–143.

Baselga, A. (2017). Partitioning abundance-based multiple-site dissimilarity into components: Balanced variation in abundance and abundance gradients. *Methods in Ecology and Evolution*, *8*, 799–808.

Harrison, S., Ross, S. J., & Lawton, J. H. (1992). Beta diversity on geographic gradients in Britain. *Journal of Animal Ecology*, *61*, 151–158.

Socolar, J. B., Gilroy, J. J., Kunin, W. E., & Edwards, D. P. (2016). How should beta-diversity inform biodiversity conservation? *Trends in Ecology & Evolution*, *31*, 67–80.

Tatsumi, S., Iritani, R., Cadotte, M.W. (2021). Temporal changes in spatial variation: partitioning the extinction and colonisation components of beta diversity. *Ecology Letters* 24, 1063–1072.

Tatsumi, S., Strengbom, J., Čugunovs, M., & Kouki, J. (2020). Partitioning the colonization and extinction components of beta diversity across disturbance gradients. *Ecology*, *101*, e03183.

### Appendix S4: Swedish fish community dataset

We retrieved the riverine fish community data from the Swedish Electrofishing Register database (Sers, 2013) via RivFishTIME (Comte et al., 2021) which is publicly available at the iDiv Biodiversity Portal (https://doi.org/10.25829/idiv.1873-10-4000). The Swedish Electrofishing Register database consists of fish community data from 33,538 electrofishing surveys in 2,992 sites across Sweden from 1951 to 2018 (as of December 19, 2021). The number of sites surveyed per year increased from 1951 to ~2005 and has remained roughly constant since then (Figure S5).

In our analyses, we used the data collected in 1990 and 2018 considering the optimal balance between the number of sites surveyed per year (i.e., the number of samples) and the length of time separating the survey years. We selected permanent sites that were surveyed in both 1990 and 2018. We excluded waterbodies in which only one site was surveyed, since we defined beta diversity as the compositional variation among multiple sites ($N \geq 2$) within each waterbody. These criteria left us with 181 permanent sites across 65 waterbodies in 1990 and 2018. There were 23 fish species found in these sites. Note that, due the intense data filtering, the results of fish community dynmics shown in this study may not represent the general trends found using the whole dataset. The impacts of local extinctions and colonizations (i.e., changes from presence to absence and vice versa) and abundance losses and gains on beta diversity of individual species are shown in Figure S6. An R script for extracting and formatting the fish community data is available at GitHub (https://github.com/communityecologist).

### References


Comte, L., Carvajal-Quintero, J., Tedesco, P.A., Giam, X., Brose, U., Erős, T...., Olden, J.D. (2021). RivFishTIME: A global database of fish time-series to study global change ecology in riverine systems. *Global Ecology Biogeography* 30, 38–50. doi:10.1111/geb.13210

Sers, B. (2013). Swedish Electrofishing RegiSter – SERS. Swedish University of Agricultural Sciences (SLU), Department of Aquatic Resources. Retrieved on December 19, 2021.


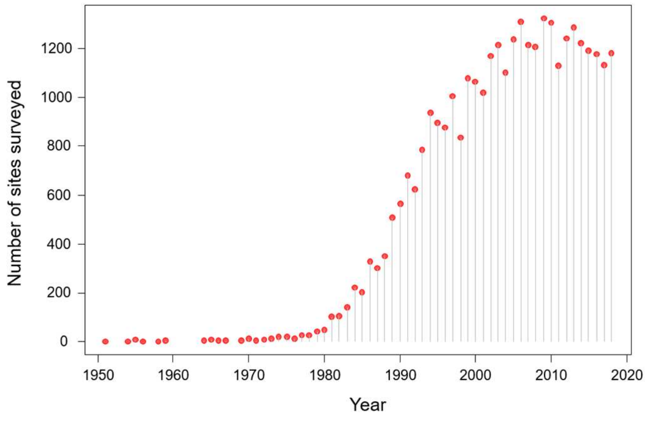


**Figure S5.** Number of sites surveyed per year in the Swedish Electrofishing Register database (Sers, 2013).

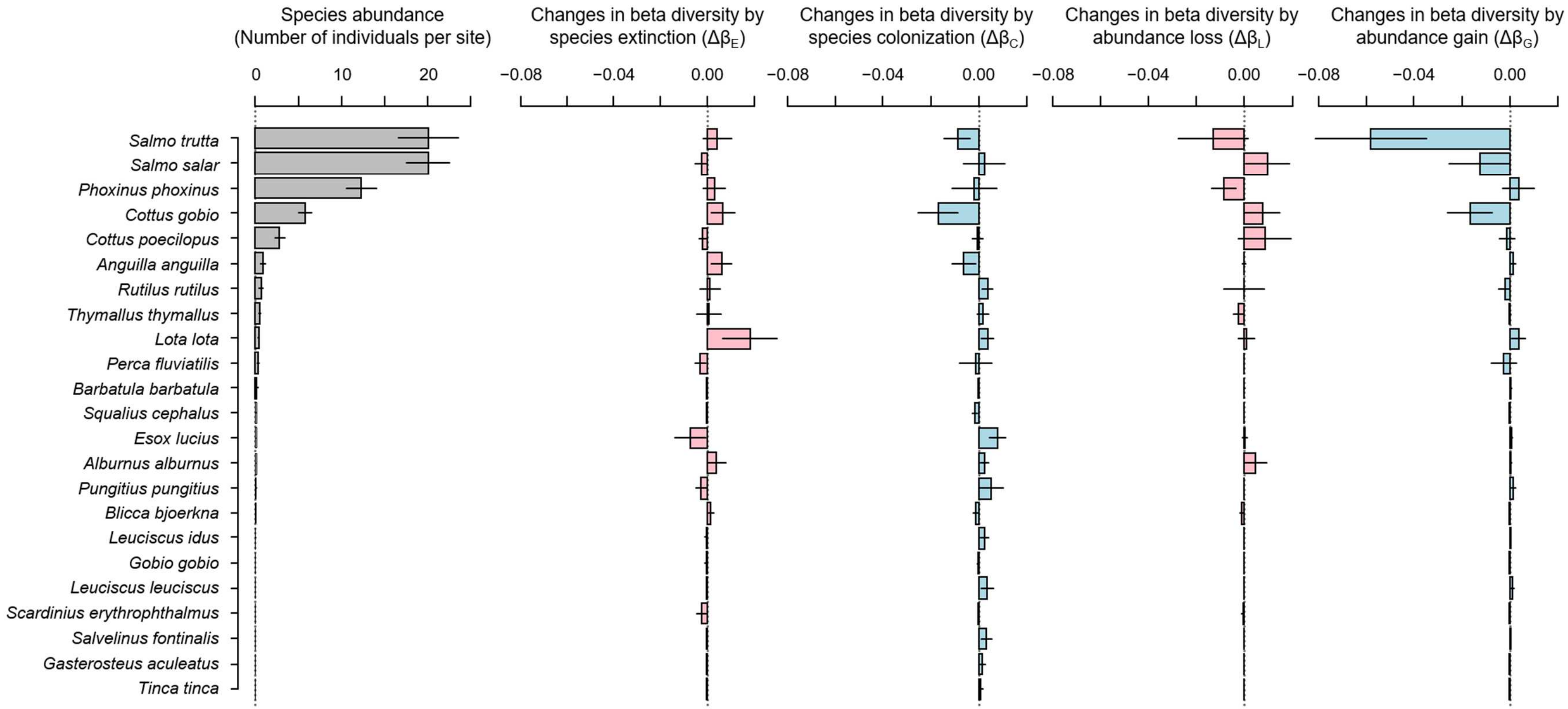


**Figure S6.** Species abundance and the beta-diversity components of riverine fish communities in 65 waterbodies across Sweden between 1990 and 2018. Beta diversity was defined as the compositional variation among multiple sampling sites within each waterbody. The red and blue bars indicate the impacts of local extinctions and colonizations (i.e., changes from presence to absence and vice versa) and abundance losses and gains on beta diversity. Bars and lines show the means ± standard errors.
